# The Drosophila MLR COMPASS-like complex regulates *bantam* miRNA expression differentially in the context of cell fate

**DOI:** 10.1101/2019.12.24.888214

**Authors:** David J. Ford, Claudia B. Zraly, John Hertenstein Perez, Andrew K. Dingwall

**Author notes:** Public Health Institute of Metropolitan Chicago, IL. Author for correspondence: Andrew K. Dingwall, PhD. Department of Cancer Biology, Stritch School of Medicine, Loyola University Chicago. Maywood, IL, USA 60153. Ph.

## Abstract

The conserved MLR COMPASS-like complexes are histone modifiers that are recruited by a variety of transcription factors to enhancer regions where they act as necessary epigenetic tools for enhancer establishment and function. A critical *in vivo* target of the Drosophila MLR complex is the *bantam* miRNA that regulates cell survival and functions in feedback regulation of cellular signaling pathways during development. We determine that loss of Drosophila MLR complex function in developing wing and eye imaginal discs results in growth and patterning defects that are sensitive to *bantam* levels. Consistent with an essential regulatory role in modulating *bantam* transcription, the MLR complex binds to tissue-specific *bantam* enhancers and contributes to fine-tuning expression levels during larval tissue development. In wing imaginal discs, the MLR complex attenuates *bantam* enhancer activity by negatively regulating expression; whereas, in differentiating eye discs, the complex exerts either positive or negative regulatory activity on *bantam* transcription depending on cell fate. Furthermore, while the MLR complex is not required to control *bantam* levels in undifferentiated eye cells anterior to the morphogenetic furrow, it serves to prepare critical enhancer control of *bantam* transcription for later regulation upon differentiation. Our investigation into the transcriptional regulation of a single target in a developmental context has provided novel insights as to how the MLR complex contributes to the precise timing of gene expression, and how the complex functions to help orchestrate the regulatory output of conserved signaling pathways during animal development.

## INTRODUCTION

COMPASS-like complexes (<u>Com</u>plex of <u>P</u>roteins <u>As</u>sociated with <u>S</u>et1) are highly-conserved chromatin modifiers responsible for methylation of the lysine 4 residue of histone 3 (H3K4), an epigenetic modification associated with active chromatin (Shilatifard, 2012). While yeast contains a single COMPASS complex, multicellular eukaryotes harbor multiple orthologous complexes specialized for specific genomic targets and methylation activity. Those containing methyltransferases KMT2C (MLL3) or KMT2D (MLL2/4) in humans and Trr in Drosophila are part of a branch of COMPASS-like complexes that we refer to as MLR (<u>ML</u>L/Trr) complexes (Fagan and Dingwall, 2019). MLR complexes are recruited to transcription enhancer regions by a variety of binding partners where they catalyze the deposition of H3K4me1, contribute to the removal of H3K27me3, and are required for recruitment of the histone acetyltransferase p300/CBP (Herz et al., 2012; Hu et al., 2013; Issaeva et al., 2007; Lai et al., 2017; Wang et al., 2016). Enhancers, also known as cis-regulatory elements, propagate transcription factor signals and control gene expression in part by forming contacts with target promoters and dramatically increasing transcription efficiency (Levine, 2010; Rickels and Shilatifard, 2018). This mechanism allows for the intricate patterns of spatiotemporal control of gene expression necessary for the normal development of most eukaryotes. Consequently, MLR complex regulation of enhancer activity is necessary for proper lineage determination, cellular differentiation, tissue patterning, and organismal development (Ang et al., 2016; Chauhan et al., 2013; Ford and Dingwall, 2015; Lee et al., 2013; Wang et al., 2016), as well as embryonic stem cell differentiation (Wang et al., 2016). The loss of MLR complex functions are causally associated with developmental disorders or lethality in multiple animal species (Andersen and Horvitz, 2007; Chauhan et al., 2012; Sedkov et al., 1999; Van Laarhoven et al., 2015). For example, germline mutations of both *KMT2C* and *KMT2D* are foundational to Kleefstra and Kabuki developmental disorders, respectively (Kleefstra et al., 2012; Ng et al., 2010); *KMT2C* and *KMT2D* are also two of the most frequently somatically mutated genes in a wide variety of human cancers, and identified as drivers of malignancy in some tumor types (Fagan and Dingwall, 2019; Ford and Dingwall, 2015). Previous work by our lab and others has determined that reduction of MLR complex activity in Drosophila larval imaginal discs results in adult organ malformation affecting tissue size and patterning. Notably, MLR complexes have been demonstrated to interact with and be necessary for proper elaboration of multiple developmental signaling pathways, and evidence suggests that alterations of Dpp/TGF-β, Hippo, and Notch signaling underlie these phenotypes (Chauhan et al., 2013; Chauhan et al., 2012; Kanda et al., 2013; Qing et al., 2014; Sedkov et al., 2003).

We sought to better understand how the MLR complex regulates critical gene enhancers and how alteration of this activity leads to disease states by examining its function in a developmental context. In Drosophila, the MLR complex contributes to the positive regulation of the expression of Tgf-β paracrine signaling molecule Dpp during wing development (Chauhan et al., 2013), and Trr physically interacts with Spen/SHARP for coactivating activity on Notch signaling targets (Oswald et al., 2016). The MLR complex also associates with the Hippo coregulator HCF and signaling effector Yorkie for proper Hippo pathway target gene activation (Nan et al., 2019; Oh et al., 2014). In the fly, these developmental signaling pathways are all linked by the miRNA *bantam*, which is a direct transcriptional target as well as a feedback regulator of these three pathways (Kane et al., 2018; Oh and Irvine, 2011; Shen et al., 2015; Wu et al., 2017). Defects resulting from alteration of these pathways can be enhanced or rescued through modulation of *bantam* levels, and *bantam* modulation alone can phenocopy these effects (Becam et al., 2011; Brennecke et al., 2003; Herranz et al., 2012; Hipfner et al., 2002; Nolo et al., 2006; Peng et al., 2009).

The *bantam* miRNA is generated from a ∼12kb non-coding precursor RNA (CR43334) and the *bantam* locus spans nearly 40kb that includes multiple tissue-specific enhancers responsible for regulating proper expression levels (Oh et al., 2013; Slattery et al., 2013). As a miRNA, *bantam* operates through translational inhibition of multiple mRNAs (Brennecke et al., 2003). It is used to control cellular function during development via regulation of targets involved in cell survival, proliferation, migration, and organ growth and patterning (Becam et al., 2011; Gerlach et al., 2019; Herranz et al., 2010; Jordan-Alvarez et al., 2017; Ma et al., 2017; Qu et al., 2017; Weng and Cohen, 2015). The best-characterized role of *bantam* is inhibition of the proapoptotic gene *hid*, regulated through Hippo signaling (Brennecke et al., 2003). The Hippo effector Yorkie (Yki) transcription factor forms heterodimers with either Scalloped (Sd) or Homothorax (Hth) to directly regulate tissue-specific *bantam* expression through multiple enhancer elements that reside up to 20 kb upstream from the CR43334 transcription unit (Slattery et al., 2013). Yki recruits the MLR complex to enhancers to activate transcription of Hippo targets, linking the MLR complex to Hippo signaling and the control of cell proliferation (Oh et al., 2014; Qing et al., 2014). We therefore reasoned that the MLR complex would likely be necessary for proper *bantam* expression and that alteration of *bantam* levels might contribute to the MLR complex loss of function phenotypes.

An ancestral genetic split of the full-length MLR methyltransferase generated separate genes in the schizophora dipterans. The Drosophila Cmi (also known as Lpt) gene is homologous to the N-terminal portion and encodes the highly conserved zinc-finger plant homeodomains (PHD) and a high mobility group (HMG) domain (Chauhan et al., 2012). Trithorax-related (Trr) contains the SET domain associated with methyltransferase activity (Sedkov et al., 2003). Despite their split into distinct genes, *Cmi* and *trr* are essential and both encoded proteins are core components of the Drosophila MLR complex (Chauhan et al., 2012). In this study we modulated the levels of Cmi and Trr in wing and eye precursor tissues, investigated *bantam*’s role in the resulting phenotypes, and assayed the function of the MLR complex on *bantam* regulation. We found that the MLR complex is bound to tissue-specific *bantam* enhancers and the complex has important functions in regulating miRNA expression, with both positive and negative effects on *bantam* transcription based on developmental context including tissue type, stage of differentiation, and cell fate. We further demonstrate that the MLR complex has critical enhancer regulatory functions within undifferentiated eye tissue cells that are required for proper *bantam* expression upon differentiation. Our results suggest that the role of MLR complexes during organismal development is more multifaceted and nuanced than previously reported.

## MATERIALS AND METHODS

### Drosophila Husbandry and Stocks

All stocks and genetic crosses were reared on standard cornmeal/dextrose medium (13% dextrose, 6% yellow cornmeal, 3% yeast extract, 1% agar, 0.3% methylparaben) at 25°C. Experimental crosses were performed at 28°C. *UAS-Cmi-IR, UAS-trr-IR*, and *UAS-Cmi* lines were previously described (Chauhan et al., 2012). The *UAS-ban-*sponge line was obtained from S. Cohen (Becam et al., 2011). The *bwe-LacZ, bee-LacZ,* and *ban*sensGFP lines were obtained from R. Mann (Slattery et al., 2013). Fly strains obtained from the Bloomington Drosophila Stock Center (BDSC) and used in this report include *Ey-Gal4* (#8220), *GawB69B-Gal4* (#1774), *DE-Gal4/TM6B* (#78371), *GMR-Gal4* (#1104), *C765-Gal4* (#36523), *en-Gal4* (#30564), and *UAS-bantam* (#60671). These strains are described in Flybase (http://flybase.bio.indiana.edu).

### Imaginal Disc Preparation, Immunofluorescence, and Imaging

Wandering third instar larvae were collected and imaginal discs dissected in ice-cold PBS, fixed in 4% formaldehyde in PBS for 15-20 minutes, then washed in PBST (PBS + 0.1% Triton X-100) three times for 5 minutes each. Washed tissues were then blocked in PBSTB (PBST + 0.1% BSA) for two hours at room temperature followed by incubation in the primary antibody diluted in PBSTB overnight at 4°C. After incubation, tissues were washed twice in PBSTB for 5 minutes each, once in PBSTB + 2% NGS for 30 minutes, and then twice more in PBSTB for 15 minutes each, followed by incubation in secondary antibody diluted in PBSTB in the dark at room temperature. Tissues were washed three times in PBST for 5 minutes each, then mounted in ProLong Gold antifade reagent with DAPI (Invitrogen).

Primary antibodies included mouse α-β-Gal (JIE7) and mouse α-Elav (9F8A9) (Developmental Studies Hybridoma Bank/Univ. of Iowa), rabbit α-GFP (GenScript) and rabbit α-Dcp-1 (Asp216) (Cell Signaling Technologies). Guinea pig α-Cmi was generated as previously described (Chauhan et al., 2012). Primary antibodies were used at 1:1000 concentration, except α-Dcp-1 was used at 1:250 concentration. Secondary antibodies were used at 1:1000 concentration and included α-Mouse, α-Rabbit, and α-Guinea Pig IgG (H+L) conjugated to Alexafluor 488 or 568 fluorophores (Life Technologies). Compound microscopy images were captured using an Olympus BX53 microscope with a Hamamatsu ORCA Flash 4.0 LT camera. Confocal microscopy images were captured using a Zeiss LSM 880 Airyscan and processed using Zeiss Zen® software. Quantification of GFP signal mean fluorescence intensity was assayed using Fiji ImageJ software to measure fluorescence intensity as mean grey value of selected areas, subtracting background (Schindelin et al., 2012).

### TUNEL Staining

TUNEL (terminal deoxynucleotidyl transferase dUTP nick end labeling) staining accomplished by collecting fixed imaginal discs and following manufacturers protocol using the In Situ Cell Death Detection Kit, Fluorescein (Roche Diagnostics). In short, fixed tissues were incubated in TUNEL solution (90% fluorescein-dUTP label solution, 10% TdT enzyme solution) for 90 minutes at 37°C in a dark humidity chamber, then washed three times in PBST for 5 minutes each and mounted for imaging.

### Scanning Electron Microscopy

Adult eyes were prepared for scanning electron microscopy using critical point drying as previously described (Wolff, 2011). SEM photography was taken at 1500X magnification using a Hitachi SU3500 microscope.

### Adult Wing Preparation, Mounting, and Imaging

Wings were dissected from adult animals and dehydrated in isopropyl alcohol for 20 minutes. After dehydration, wings were mounted in DPX mountant (Fluka). Images were captured using a Leica MZ16 microscope with Leica DFC480 camera.

### ChIP-seq

Chromatin from whole animals was collected, prepared, and analyzed according to (Zraly et al., 2020).

### miRNA extraction, cDNA synthesis and qRT-PCR

Wing discs from 50 Drosophila third instar larvae of the appropriate genotype were dissected and miRNA was prepared using the miRVana isolation kit (Life technologies) to enrich for small RNAs, according to manufacturer’s protocols. RNA (10ng) was reverse transcribed using Multiscribe reverse transcriptase (ThermoFisher Scientific) and TaqMan small RNA assay RT primers specific for bantam (Assay ID: 000331) and 2S RNA (Assay ID: 001766; control), according to manufacturer’s protocols. For qRT-PCR of miRNAs, TaqMan Universal PCR Master Mix II (No UNG, ThermoFisher Scientific) and miRNA-specific TaqMan microRNA Assay was used. To enrich for long RNAs, total RNA was also isolated using the miRVana kit according to manufacturer’s protocols. The RNA was DNase digested, and reverse transcribed, as previously described (Chauhan et al., 2013). The primer sequences used to amplify *bantam* precursor, CR43334 are as follows: Forward primer: 5’-GCGATGTATGCGTGTAGTTAAAG-3’; Reverse primer: 5’-CCACTTTGTCGATCGTTTCATG-3’. Real time qPCR was performed on a QuantStudio6 Flex Real Time PCR System (BioRad). The 2^−ΔΔ^*^C^*^T^ method was used for quantification.

### Statistical Analysis

Significant difference of eye phenotype severity between genetic populations was measured using Pearson’s Chi-Squared Test. Significant difference of mean fluorescence intensity in eye and wing discs was measured using Student’s T-test.

## RESULTS

### The MLR Complex is Necessary for Suppressing *bantam* Expression in the Developing Wing

We previously reported that modulation of Cmi levels during development leads to a variety of defects, including wing vein pattern disruptions, small and rough eyes, decreased expression of ecdysone hormone regulated genes and effects on organismal growth (Chauhan et al., 2012). In the developing wing, reduced *Cmi* function leads to retraction of the L2 and L5 longitudinal veins and posterior crossvein, while overexpression causes smaller wings with bifurcation of distal veins, as well as ectopic vein formation. These phenotypes result in part from alteration of Dpp/Tgf-β signaling (Chauhan et al., 2013). *Cmi* or *trr* knockdown in larval eye discs cause reduction in eye size, while overgrowth is observed at low penetrance with Cmi overexpression (Chauhan et al., 2012; Sedkov et al., 2003). The *Cmi* and *trr* loss of function phenotypes likely reflect disruptions in gene regulatory networks that control development in response to multiple signaling pathways due to improper enhancer regulation. Indeed, the MLR complex has been implicated in regulating aspects of the Hippo and Notch pathways as well (Kanda et al., 2013; Oh et al., 2014; Qing et al., 2014).

The Tgf-β, Hippo and Notch pathways share a common transcriptional target, the *bantam* miRNA, and regulation of *bantam* expression by these pathways is necessary for proper imaginal disc formation (Attisano and Wrana, 2013; Becam et al., 2011; Boulan et al., 2013; Doumpas et al., 2013; Herranz et al., 2012; Kane et al., 2018; Li and Padgett, 2012; Martin et al., 2004; Oh and Irvine, 2011; Oh et al., 2014; Qing et al., 2014; Wu et al., 2017; Zhang et al., 2013). Given the requirement for the MLR complex in regulating targets of these pathways, we hypothesized that the *bantam* miRNA is a potential direct target of the complex. To test this, we turned to the wing imaginal disc, which broadly expresses *bantam* in a stable pattern (Brennecke et al., 2003). To reduce MLR complex activity, we used Gal4-driven RNAi to knockdown either *Cmi* or *trr* in the wing disc. Cmi and Trr were chosen because they are core subunits necessary for function of the MLR complex (Shilatifard, 2012; Zraly et al., 2020). If *bantam* is a regulatory target of the complex, then *bantam* expression should be sensitive to a reduction in MLR activity. Surprisingly, we observed increased transcript levels of both *bantam* miRNA and its precursor transcript lncRNA:CR43334 upon knockdown of *Cmi* or *trr* in the wing disc (*C765-Gal4>Cmi-IR* and *C765-Gal4>trr-IR*) compared to wild type (Fig. 1A), suggesting that *bantam* expression was negatively regulated or repressed by the MLR complex in this tissue. We verified these results using a *bantam* sensor GFP (*ban*sensGFP) construct that functions as a constitutively expressed inverse reporter of *bantam* activity, such that the higher the expression of *bantam* in a cell, the lower the levels of GFP (Brennecke et al., 2003). Compared to control wing discs and wild type anterior tissue, knockdown of *Cmi* or *trr* within the posterior wing compartment using the e*n-Gal4* driver results in decreased GFP reporter expression, reflecting increases in *bantam* levels in the developing wing (Fig. 1B). These data demonstrate that the MLR complex plays a necessary role in attenuating *bantam* expression in the developing wing.

**Figure 1.**
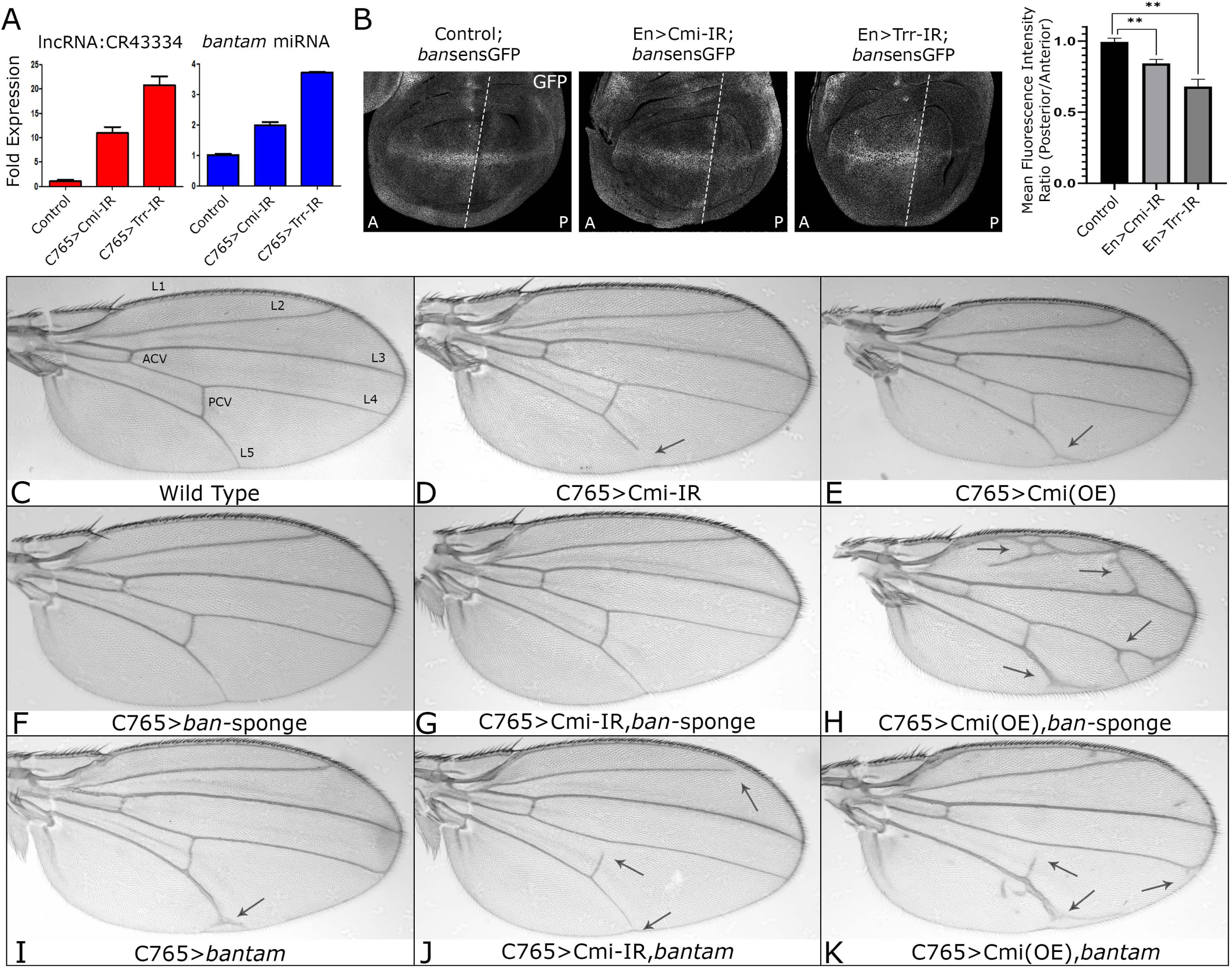
*The MLR complex genetically interacts with* bantam *during wing development.* (A) RT-qPCR was performed on whole wing discs. Both the *bantam* precursor lncRNA:CR43334 and processed *bantam* miRNA are upregulated upon *Cmi* or *trr* knockdown. (B) *en-Gal4* was used to knockdown *Cmi* or *trr* in the posterior wing disc; a *bantam*-GFP inverse sensor construct was used to compare *bantam* miRNA levels between posterior (P) and anterior (A) of organ. (Left) Tissue in the posterior of *en>Cmi-IR* and *en>trr-IR* organs demonstrate increased *bantam* levels (lower GFP). (Right) posterior/anterior ratio of *bantam* sensor signal quantified; ** = p ≤ 0.01. (C-K) *C765-Gal4* was used to drive expression of genetic constructs modulating levels of Cmi or *bantam* in the wing disc. Compared to wild type wings (C), knockdown of *Cmi* (Cmi-IR) (D) causes retraction of wing veins, most commonly L5. (E) Cmi overexpression results in smaller wings and ectopic distal veins. (F) Reduction of *bantam* activity via expression of a *bantam*-specific miRNA sponge (UAS-*ban-*sponge) does not alter adult wing phenotype. (G) *bantam* reduction in the background of *Cmi* knockdown completely rescues the wild type phenotype. (H) *bantam* reduction alongside Cmi overexpression enhances the phenotype, causing ectopic vein formation. (I) Overexpressing *bantam* during wing formation causes end vein forking of L5. (J) *bantam* overexpression in parallel with *Cmi* knockdown enhances the Cmi loss-of-function phenotype. (K) Overexpression of both Cmi and *bantam* results in enhanced vein forking, crossvein retraction, and rescues wing size. Black arrows highlight vein formation defects.

As *bantam* acts as a negative feedback regulator of Dpp signaling in the wing disc (Kane et al., 2018), we investigated whether the *Cmi* reduced function wing phenotypes (Chauhan et al., 2013; Chauhan et al., 2012) (Fig. 1D-E) were also sensitive to *bantam* levels. Genetic interaction tests between *Cmi* and *bantam* were performed using the Gal4-UAS system (Brand et al., 1994) expressing transgenic shRNAi or overexpression constructs to modulate Cmi levels in the entire larval wing disc; in parallel, *bantam* activity was either increased through overexpression or depleted using a *bantam*-specific miRNA sponge (*ban-*sponge) (Herranz et al., 2012) (Fig. 1 C-K, Table 1). Simultaneous knockdown of *Cmi* and reduction of *bantam* activity (*C765-Gal4>Cmi-IR,* ban*-sponge*) results in phenotypically wild type wings, demonstrating complete suppression of the short vein phenotype associated with reduced *Cmi* function (Fig. 1G). Conversely, overexpression of *bantam* combined with *Cmi* knockdown results in greater retraction of L2 and L5 veins, thus enhancing the severity of the *Cmi* loss of function phenotype (Fig. 1J). In the background of *Cmi* overexpression (*C765-Gal4>Cmi(OE)*), *bantam* activity depletion leads to an increase in the severity of vein bifurcation and ectopic vein formation (Fig. 1H). Concurrent overexpression of *Cmi* and *bantam* does not appear to suppress the *Cmi* gain-of-function vein phenotype, although wing size is increased to match wild type (Fig. 1K). These data support a hypothesis that the MLR complex and *bantam* interact genetically. Additionally, the inverse relationship between MLR activity and *bantam* expression suggests that the role of MLR in suppressing *bantam* levels may be mechanistically involved in the dose dependent *Cmi* wing phenotypes.

**Table 1.**
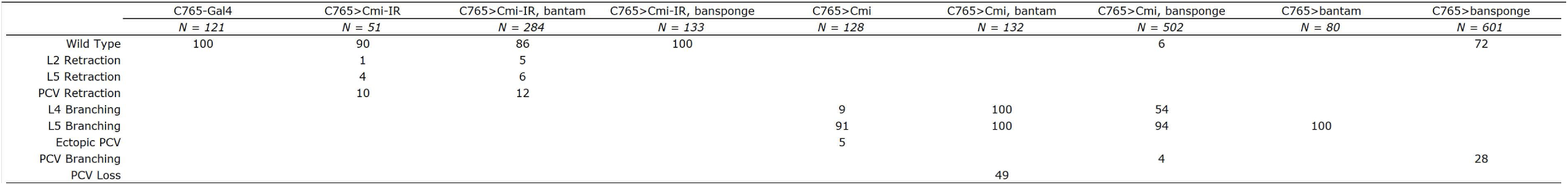
Wing phenotype scoring for Cmi/bantam genetic interaction.

### The MLR Complex Directly Modulates Tissue-Specific *bantam* Enhancers

The reduction of *bantam* levels in wing discs in response to *Cmi* or *trr* knockdown may be the result of direct or indirect regulatory mechanisms. If the MLR complex directly regulates the expression of *bantam*, it would do so through control of *bantam*-specific enhancers. Such regulatory regions have been identified as necessary and sufficient for transcription factor control of *bantam* locus expression (Oh and Irvine, 2011; Slattery et al., 2013). A *bantam* wing enhancer locus was previously identified as a cis-regulatory element residing approximately 20kb upstream of the lncRNA:CR43334 transcription start site (Fig. 2A); an enhancer reporter of this region recapitulates the expression pattern of the *bantam* miRNA in the wing imaginal discs, demonstrating tissue-specific regulatory activity (Slattery et al., 2013). To identify MLR complex-bound regions of the genome, we recently performed Cmi ChIP-sequencing at various developmental stages (Zraly et al., 2020). Examination of Cmi enrichments near *bantam* revealed localization throughout the locus of the precursor RNA as well as the upstream regulatory region, including peaks at the identified wing enhancer (Fig. 2A). Although Cmi is broadly enriched at the bantam locus at multiple stages of development it is only present at the wing enhancer at the wandering third instar and prepupal stages. To determine if these binding events are associated with regulatory activity by the MLR complex, we took advantage of a LacZ reporter construct created to verify the activity patterns of these enhancers (Slattery et al., 2013). Knockdown of either *Cmi* or *trr* in the wing lead to increased activity of the *bantam* wing enhancer in the cells normally expressing the miRNA (Fig. 2B), corresponding with the previous genetic and biochemical data in the wing (Fig. 1) and demonstrating that the MLR complex is critical for attenuating *bantam* expression during wing development through restraints on enhancer activity.

**Figure 2.**
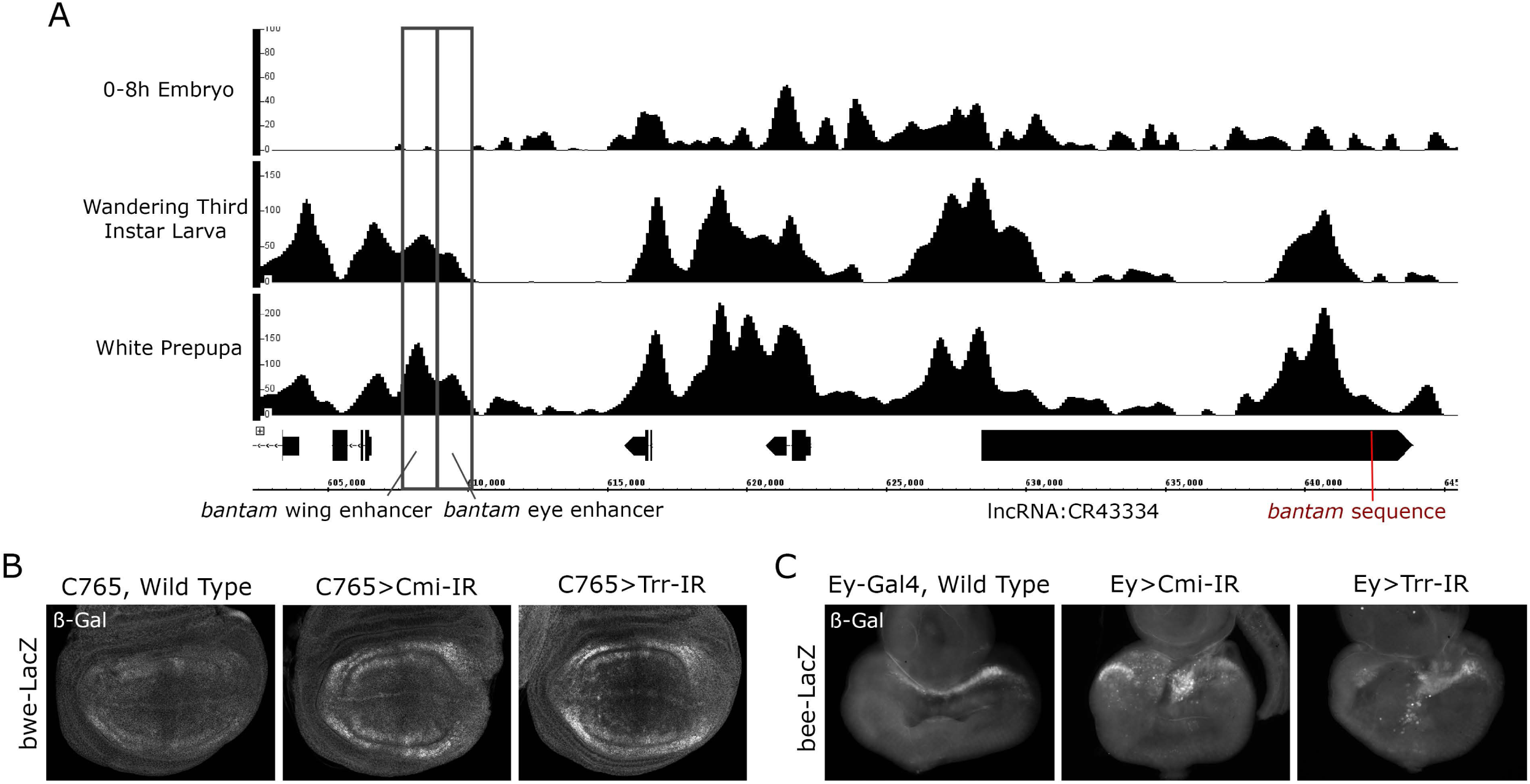
*The MLR complex localizes to and regulates tissue-specific bantam enhancers.* (A) ChIP-seq was used to assay binding of MLR subunit Cmi across the *bantam*-containing lncRNA locus as well as at upstream tissue-specific enhancer regions. Binding profiles across different developmental timepoints including early embryo (E0-8), wandering third-instar larva (W3L), and white prepupa (WPP), demonstrate consistent localization of the complex to the lncRNA locus as well as previously identified eye- and wing-specific enhancer regions (Slattery et al., 2013) during imaginal disc development. (B) *Cmi* or *trr* was knocked down throughout the wing disc using *C765-Gal4*; a *bantam* wing enhancer-β-Gal reporter *(bwe-LacZ)* was used to compare enhancer activity levels. *C765>Cmi-IR* and *C765>trr-IR* discs demonstrate increased *bantam* wing enhancer activity levels compared to control. (C) *Cmi* or *trr* was knocked down throughout the eye pouch of the eye disc using *Ey-Gal4*. A *bantam* eye enhancer-β-Gal reporter *(bee-LacZ)* was used to compare enhancer activity levels. *Ey>Cmi-IR* and *Ey>trr-IR* organs demonstrated disrupted *bantam* eye enhancer activity patterns compared to control.

Our ChIP-seq data also revealed developmentally-associated Cmi enrichment at an eye disc-specific *bantam* enhancer (Fig. 2A) (Slattery et al., 2013), suggesting that the MLR complex also plays a role in *bantam* regulation in the developing eye. An enhancer-LacZ reporter of this region demonstrates activity at the very anterior margin of the eye pouch bordering the antennal section (Fig. 2C). Rather than a clear increase or decrease in activity upon *Cmi* or *trr* knockdown, this stable pattern is disrupted with sporadic sections lacking enhancer activity and apparent ectopic activation in other sections. These data support a hypothesis that the MLR complex also directly regulates the *bantam* eye enhancer, but its role may be in modulating enhancer activity and restricting activation to certain cells.

### The MLR Complex Interacts with *bantam* During Eye Development

To investigate potential MLR complex regulation of *bantam* during eye development, we first turned to previously identified eye phenotypes. Reduced MLR complex function in the Drosophila eye imaginal disc results in rough and shrunken adult eyes, including disorganized ommatidia (Chauhan et al., 2012; Kanda et al., 2013; Sedkov et al., 2003). If this effect is sensitive to *bantam* levels in the eye, it would suggest mechanistic interaction between the miRNA and the complex during eye development. *Eyeless-Gal4* (Ey-Gal4) was used to express *Cmi-* or *trr-*specific shRNAi constructs in the entire eye pouch of the eye-antennal imaginal disc (*Ey-Gal4>Cmi-IR* and *Ey-Gal4>trr-IR*), resulting in phenotypes ranging in severity from small eyes with slight roughness on the posterior margin to near-complete loss of eye tissue (Fig. 3A). SEM imaging of adult eyes of both knockdown phenotypes revealed disruptions in ommatidial patterning accompanied by bristle loss and duplication, as well as lens fusion (Fig. 3D-E). Phenotypes associated with *trr* knockdown demonstrated higher penetrance and expressivity compared to *Cmi* knockdown (Fig. 3B,D-E). This difference is consistent with the fact that the MLR complex is able to form in the absence of Cmi, whereas loss of Trr causes instability of the entire complex (Zraly et al., 2020). The variable expressivity of the *Cmi* knockdown phenotype is more sensitive, and therefore allows us to more clearly observe changes in phenotype severity due to modulation of *bantam* levels. *Ey-Gal4>bantam* overexpression in the eye disc results in similar rough and shrunken eyes (Fig. 3B,F), and *bantam* overexpression concurrent with *Cmi* or *trr* knockdown significantly enhances the rough and shrunken phenotype (Fig. 2B,G-H). Previous studies in which *bantam* was overexpressed exclusively within the differentiating eye have reported similar disruptions of ommatidial patterning due to excess interommatidial cells (Nolo et al., 2006; Thompson and Cohen, 2006). Reduction of *bantam* activity via *ban-*sponge expression suppresses the *Cmi* knockdown-associated defects despite the fact that *bantam* depletion alone does not produce a phenotype (Fig. 2B,I-K). These data reinforce that the MLR complex has a role in controlling *bantam* levels during eye development.

**Figure 3.**
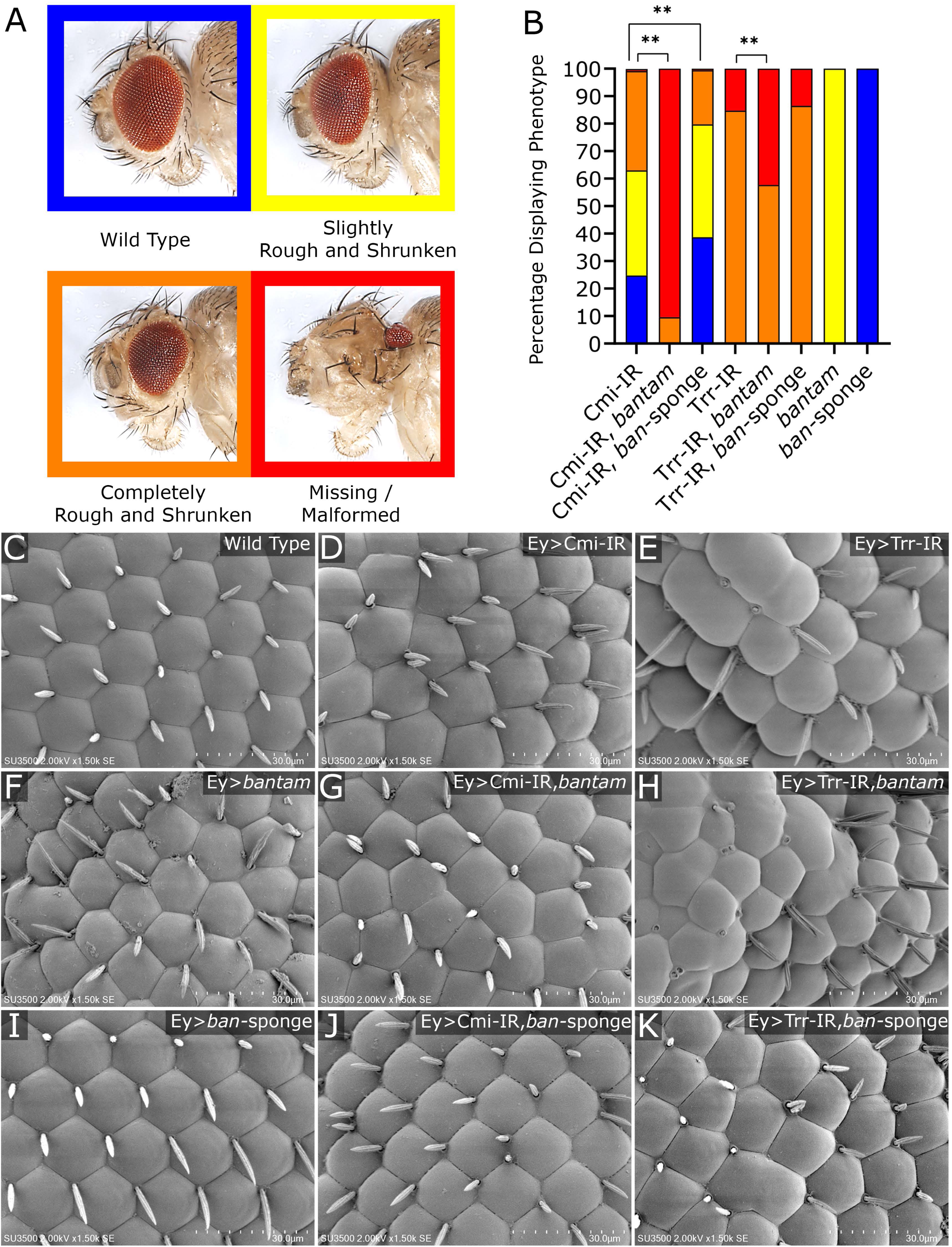
*The MLR complex genetically interacts with* bantam *during eye development. Ey-Gal4* was used to drive expression of genetic constructs modulating levels of Cmi, Trr, or *bantam* in the eye disc. (A) Knockdown of *Cmi* or *trr* during eye development results in rough and shrunken eyes that range in penetrance and expressivity. (B) Adults of the listed genotypes were scored according to eye phenotype severity; N > 50 for all genotypes; ** = p ≤ 0.01. Overexpression of *bantam* significantly enhances the *Cmi* or *trr* knockdown phenotypes, while reduction of *bantam* activity using the *ban*-sponge suppresses the *Cmi* knockdown phenotype. Overexpression of *bantam* alone causes slightly rough and shrunken eyes; expression of the *ban*-sponge alone has no phenotypic effect. (C-K) SEM images of adult compound eyes demonstrate that roughness is due to ommatidial patterning defects. As compared to wild type ommatidia (C), *Cmi* knockdown (D) and *trr* knockdown (E) eyes display ommatidial crowding, lens fusion, and bristle loss and duplication. This is also seen when *bantam* is overexpressed alone (F), in the Cmi-IR background (G), and the Trr-IR background (H). (I) Reduction of *bantam* activity has no effect on ommatidial patterning and appears to suppress the effect of *Cmi* (J) or *trr* (K) knockdown.

### The MLR Complex Regulates Apoptosis in the Developing Eye

The reduced eye size and disrupted ommatidia phenotype caused by knockdown of *Cmi* or *trr* may reflect elevated apoptosis, decreased proliferation, and/or defects in cellular differentiation. Clones of *trr* mutant cells in the eye display small patches of Caspase-3 positive staining (Kanda et al., 2013), implicating a role for the MLR complex in cell survival. A well-characterized function of *bantam* in the eye disc is the inhibition of apoptosis through translational blocking of proapoptotic *hid* (Brennecke et al., 2003; Grether et al., 1995). Although the genetic data suggest that increased *bantam* function is associated with the rough and shrunken phenotype, it is possible that dysregulation of the miRNA may alter cell survival through other means in *Cmi/trr* knockdown eyes. If loss of MLR activity does in fact deleteriously affect cell survival, this would occur in the eye imaginal disc. The eye disc is a useful developmental model as, unlike the wing, it contains cells at multiple stages of differentiation and development. A mobile boundary known as the morphogenetic furrow induces differentiation as it migrates from the posterior to the anterior margin of the eye pouch, marking the induction of differentiation into separate lineages of proneuronal and interommatidial cells (Fig. 5A). Therefore, tissue anterior to the furrow remains undifferentiated while that posterior to the furrow has begun synchronized differentiation, eventually giving rise to the multiple cell types that will make up the adult compound eye (Cagan, 2009). To address whether elevated apoptosis is causal to the eye phenotype, we utilized TUNEL staining. Eye discs undergo sporadic apoptosis in undifferentiated tissue and regulated pruning of cells once the morphogenetic furrow passes and synchronized differentiation commences (Brachmann and Cagan, 2003), yet knockdown of either *Cmi* or *trr* in the eye pouch results in high levels of TUNEL staining in the anterior undifferentiated section of the eye pouch, with a significant concentration of these cells on the dorsal-ventral midline (Fig. 4A), the same location identified in *trr* mutant clones (Kanda et al., 2013). To verify caspase cascade and test cell autonomy of the effect, we knocked down *Cmi* or *trr* exclusively in the dorsal eye region via the *DE-Gal4* driver and stained eye discs for presence of activated effector caspase Dcp-1 (Song et al., 1997) (Fig. 4B). These results demonstrate that caspases are activated solely within *Cmi*/*trr* knockdown cells, suggesting that reduction of MLR activity in undifferentiated eye cells has a deleterious effect on the survival of those cells. (Fig. 4B). To investigate a potential role of *bantam*, Dcp-1 activation was examined in discs overexpressing *bantam* or the *ban-*sponge in combination with *Cmi* or *trr* knockdown and compared to control discs (Fig. 4C-E). While the role of *bantam* and the effects of overexpression within differentiating eye cells has been described (Thompson and Cohen, 2006), the effects of *bantam* modulation in undifferentiated eye tissue anterior to the morphogenetic furrow is uncertain. Surprisingly, overexpression of the miRNA within the entire eye pouch induces widespread caspase activation in undifferentiated cells, causes overgrowth of the eye pouch, and enhances the Dcp-1 activation phenotype in a *Cmi* or *trr* knockdown background (Fig. 4F-H). Reduction in *bantam* activity through the *ban*-sponge suppresses this phenotype (Fig. 4I-K).

**Figure 4.**
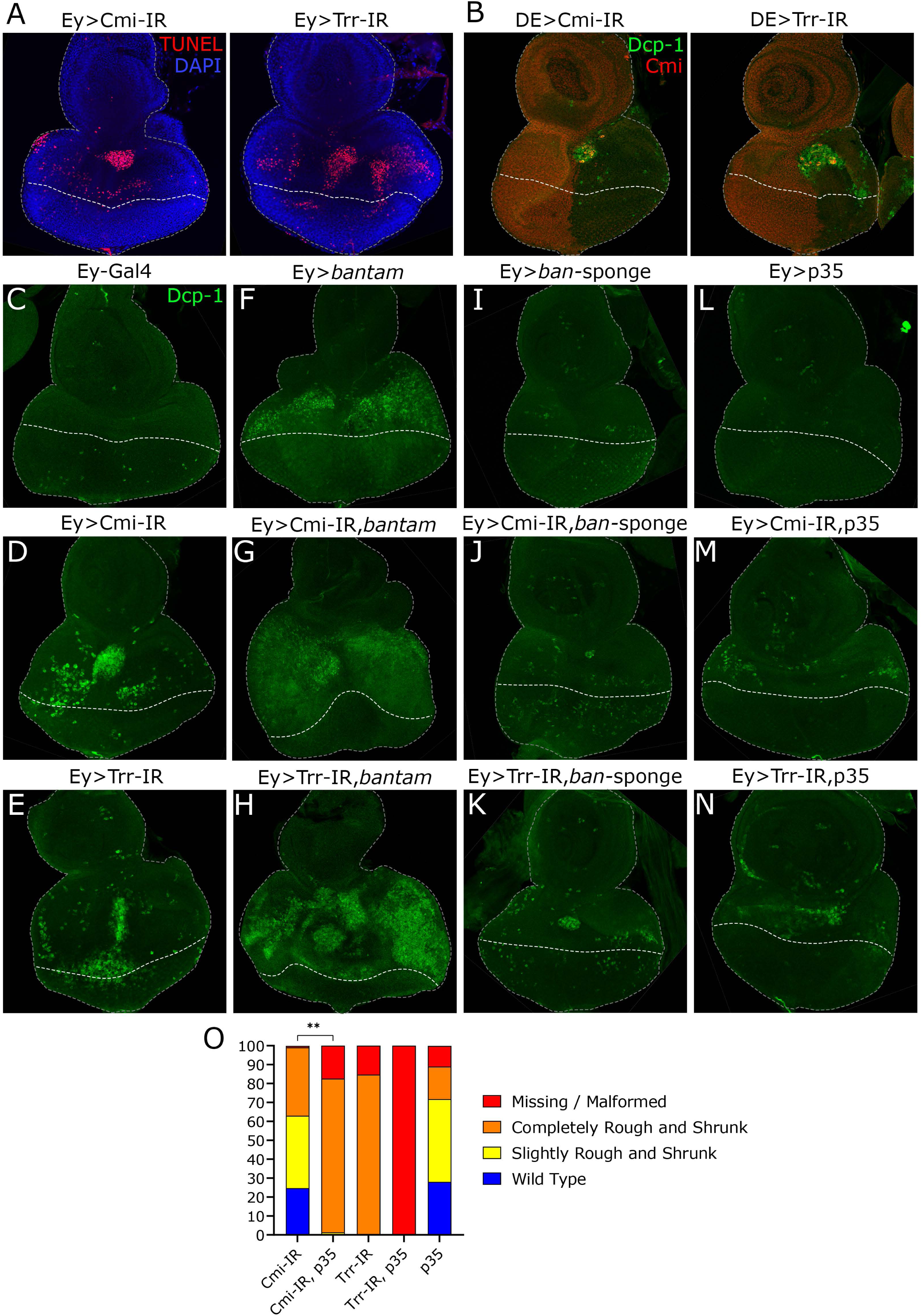
*The MLR complex suppresses caspase activation in the undifferentiated eye.* Eye disc were dissected from wandering third instar larvae and assayed for apoptosis; the morphogenetic furrow is marked by a thick dotted line and the discs are outlined by a thin dotted line. (A) *Cmi* or *trr* was knocked down in the entire developing eye pouch using *Ey-Gal4*. TUNEL staining (red) displays apoptotic cell death concentrated in a cluster of apoptotic cells on the dorsal-ventral midline anterior to the morphogenetic furrow. (B) *Cmi* or *trr* was knocked down in the dorsal half of the eye pouch using *DE-Gal4*. Knockdown is visualized by staining for Cmi (red), which is reduced upon knockdown of either *Cmi*/*trr*. Cleaved Dcp-1 staining (green) demonstrates that caspase cascade is induced only within tissue knocking own *Cmi*/*trr* and not within neighboring cells. (C-N) *Ey-Gal4* was used to drive expression of genetic constructs modulating levels of Cmi, Trr, or *bantam* in the eye disc. Apoptosis in the eye disc was assayed by staining for cleaved effector caspase Dcp-1 (green). (C) Wild type eye discs demonstrate low sporadic caspase activation. (D-E) Knockdown of *Cmi* or *trr* in the eye pouch using *Ey-Gal4* results in increased caspase activity in undifferentiated cells concentrated in a cluster of cells on the dorsal-ventral midline anterior to the morphogenetic furrow. (F) Overexpression of *bantam* causes eye pouch overgrowth and caspase activation in undifferentiated tissue. (G-H) Increased *bantam* expression in the background of *Cmi*/*trr* knockdown enhances caspase activation. While reduction of *bantam* activity through expression of the *ban-*sponge has no effect on caspase activation (I), it suppresses the apoptotic induction of *Cmi*/*trr* knockdown organs (I-K). (L) Caspase inhibitor p35 represses caspase activation. (M-N) p35 expression alongside *Cmi*/*trr* knockdown suppresses the caspase phenotype. (O) Adult eye phenotype was scored and quantified. p35 expression phenocopies the rough and shrunken phenotype of *Cmi*/*trr* knockdown and enhances the phenotype in a *Cmi*/*trr* knockdown background; ** = *p*≤0.01. Simultaneous *trr* knockdown and p35 expression results in synthetic lethality; therefore, statistical significance between Trr-IR and Trr-IR, p35 not achieved due to low number of surviving adults.

**Figure 5.**
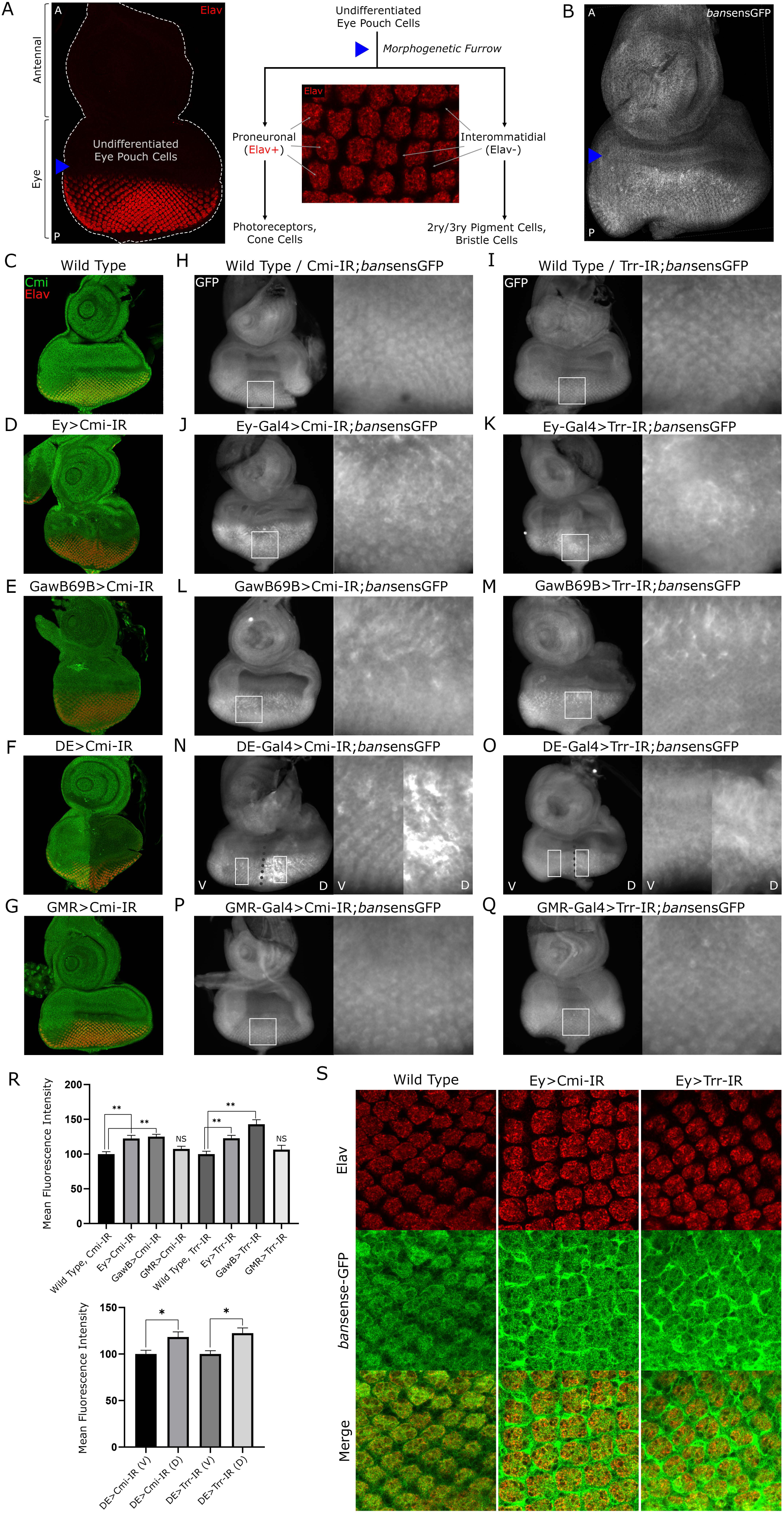
*The MLR complex regulates* bantam *expression.* (A) Diagram of compound eye differentiation in the W3L eye disc. To the left, the eye disc is outline in a thin dotted line. It is comprised of an anterior (A) antennal pouch and a posterior (P) eye pouch. The eye pouch is separated into anterior undifferentiated cells and posterior differentiating cells by the morphogenetic furrow (marked by blue arrow). After passage of the furrow, a cell fate decision is made between proneuronal lineage, positive for Elav (red), and interommatidial lineage. During metamorphic compound eye development, the proneuronal lineage gives rise to photoreceptors and cone cells while the interommatidial lineage gives rise to secondary and tertiary pigment cells as well as bristle cells. (B) Example of *ban*sensGFP expression in a wild type eye disc. Undifferentiated eye cells anterior to the morphogenetic furrow (marked by blue arrow) have relatively high *bantam* levels (low *ban*sensGFP), and differentiating cells posterior to the furrow have relatively low *bantam* levels high *ban*sensGFP). (C-G) *Cmi* was knocked down using various Gal4 drivers to visualize driver expression pattern. Expression pattern of these drivers is visualized by immunostaining Cmi (green) and Elav (red), which labels proneuronal cells posterior to the morphogenetic furrow. (D) *Ey-Gal4* drives knockdown within the eye pouch. (E) *GawB69B-Gal4* drives ubiquitously throughout the organ. (F) *DE-Gal4* drives only within the dorsal half of the eye pouch. (G) *GMR-Gal4* drives only posterior to the morphogenetic furrow. (H-Q) The *bantam*-GFP inverse sensor construct (*ban*sensGFP) was used to assay *bantam* miRNA levels. Magnified views of developing ommatidia posterior to the furrow are displayed to the right of each eye disc. *bantam* levels in the undifferentiated anterior tissue anterior to the furrow remain unchanged in all genotypes. (H-I) In control discs, bantam levels are relatively high anterior to the furrow (low GFP) and lower posterior (high GFP). (J-O) When *Cmi* or *trr* is knocked down both anterior and posterior to the furrow (*Ey>*, *GawB69B>*, and *DE>*), *bantam* levels appear to decrease in the differentiating posterior tissue. (P-Q) If Cmi/Trr are lost only within this differentiating tissue (*GMR>*), *bantam* levels remain unchanged compared to control. (R) Mean fluorescence intensity anterior to the furrow was quantified from cohorts of each genotype; N ≥ 10 for all genotypes; * = p≤0.05, ** = p≤0.01, NS = not significant. (S) Developing ommatidia in eye discs were stained for pro-neuronal marker Elav (red) and *ban*sensGFP (green). Control organs demonstrate colocalization of GFP and Elav. Upon knockdown of either *Cmi* or *trr* (Ey>), changes in *bantam* expression vary by cell fate. In proneuronal cells *bantam* levels are increased (lower GFP), and in interommatidial cells bantam levels are decreased (higher GFP).

Given the fact that caspase activation in undifferentiated tissue as well as rough and shrunken adult eyes are both associated with reduced MLR complex activity and are similarly sensitive to *bantam* levels, we hypothesized that the aberrant cell death in the undifferentiated eye is causal to smaller adult organs with patterning defects. To verify this, caspase activation in the eye disc was suppressed by expression of p35, a baculovirus substrate inhibitor of caspases including Dcp-1 (Song et al., 2000; Zoog et al., 1999). If increased cell death in undifferentiated eye tissue is mechanistically linked to the rough and shrunken adult eyes, then suppression of apoptosis by p35 should rescue the adult phenotype. Intriguingly, while p35 successfully reduces apoptotic activation by *Cmi* or *trr* knockdown (Fig. 4L-N), it significantly enhances the adult phenotypes (Fig. 4O). From these data, we conclude that the adult eye phenotype is not a result of the induction of apoptosis in the undifferentiated eye disc. Rather, these are two independent effects both resulting from the loss of MLR complex activity.

### The MLR Complex Regulates *bantam* Expression in a Tissue- and Differentiation-Specific Context

The phenotypes caused by MLR subunit knockdown both in the eye and in the wing are similarly sensitive to *bantam* levels, and the MLR complex directly regulates both the eye- and wing-specific *bantam* enhancers, suggesting that the MLR complex attenuates *bantam* expression in the eye imaginal disc as it does in the wing disc. To investigate this, the *ban*sensGFP construct was used as an inverse assay of *bantam* levels in the eye disc. In wild type eye discs, *bantam* levels remain high in undifferentiated tissue and vary by cell type once differentiation commences (Fig. 5B) (Tanaka-Matakatsu et al., 2009). Knockdown of *Cmi* or *trr* driven by *Ey-Gal4* (which results in caspase activation and malformed adult eyes) occurs within both undifferentiated and differentiating eye tissue (Fig. 5D). As loss of MLR activity induces apoptosis in undifferentiated tissue anterior to the furrow, we anticipated altered *bantam* expression in this region. Unexpectedly, *bantam* levels in cells anterior to the morphogenetic furrow were unchanged compared to control (Fig. 5J-K). Posterior to the furrow, however, *ban*sensGFP signal significantly increases, representing a decrease in *bantam* expression. To verify these results, two additional Gal4 drivers were used: *GawB69B-Gal4* drives expression ubiquitously throughout the entire disc (Fig. 5E); *DE-Gal4* drives expression within the dorsal half of the eye pouch, leaving the ventral half as an internal wild type control (Fig. 5F). Knockdown of either *Cmi* or *trr* in these undifferentiated eye cells does not alter *bantam* expression in those cells (Fig. 5L-O), suggesting that the caspase activation effect is not caused by dysregulated *bantam* expression. Also, in each case of *Cmi*/*trr* knockdown in both undifferentiated and differentiating cells, *ban*sensGFP signal is significantly increased in those differentiating (Fig. 5J-O,R). These results suggest that the MLR complex is not required for maintaining *bantam* transcription in undifferentiated eye disc cells but may have critical functions in epigenetic reprogramming control of cell-type specific *bantam* enhancers during cell fate transitions. This was tested using *GMR-Gal4*, which drives knockdown only in those differentiating cells posterior to the furrow (Fig. 5G). Unexpectedly, knockdown of *Cmi*/*trr* only in this region had no effect, resulting in eye discs demonstrating wild type expression of *bantam* (Fig. 5P-Q,R). Taken together, these data suggest that the MLR complex has regulatory function in undifferentiated cells necessary for proper *bantam* transcription upon differentiation but is dispensable for maintaining *bantam* expression programming once differentiation has commenced.

The downregulation of *bantam* expression observed in differentiating cells did not appear to be uniform; rather, *bantam* expression appeared to vary by cell type in response to knockdown of *Cmi* or *trr* (Fig. 3H-M). These cells have begun cell fate decisions that separate them into patterns of specific lineages (Fig. 5A), suggesting that cells of only particular lineages are affected by MLR subunit knockdown. In wild type eye discs, *bantam* is downregulated in proneuronal cells at the center of each developing ommatidia and upregulated in the interommatidial cells bordering the compound eye units (Fig. 5S) (Tanaka-Matakatsu et al., 2009). Differentiating cells in *Cmi* or *trr* knockdown discs were closely examined via confocal microscopy, revealing two separate effects: *bantam* is simultaneously upregulated in the differentiating proneuronal ommatidial cells (marked by Elav staining) and downregulated in interommatidial cells. These data reveal not only that the MLR complex has an important and unexpected role in establishing *bantam* expression levels prospectively in undifferentiated eye cells, but also that this regulatory activity is necessary for proper cell type-specific *bantam* expression in subsequent cell generations in response to differentiation signals.

## DISCUSSION

The highly conserved MLR COMPASS complexes serve to commission, bookmark and maintain epigenetic *cis*-regulatory controls on transcription enhancers, thus helping to fine tune signaling pathways that are critical for animal development and cellular homeostasis (Fagan and Dingwall, 2019; Rickels et al., 2017; Wang et al., 2018; Zraly et al., 2020). We recently showed that the complex displayed developmental context-dependent chromatin enrichments that linked regulatory functions of the complex to potential *in vivo* target genes, including those important for developmental pattern formation and growth control (Zraly et al., 2020). We therefore sought to explore *in vivo* mechanisms of target gene regulation by the Drosophila MLR complex and discovered two key features: 1) We have identified the miRNA *bantam* as an important regulatory target of the complex during tissue development with roles in both activating and repressing *bantam* expression depending on the context of cell fate (Fig. 6). To our knowledge, this is the first demonstration that the complex has a role in both positive and negative regulation of a single transcriptional target in the same tissue. 2) The MLR complex has important functions in enhancer maintenance and sustaining cell survival in undifferentiated cells.

**Figure 6.**
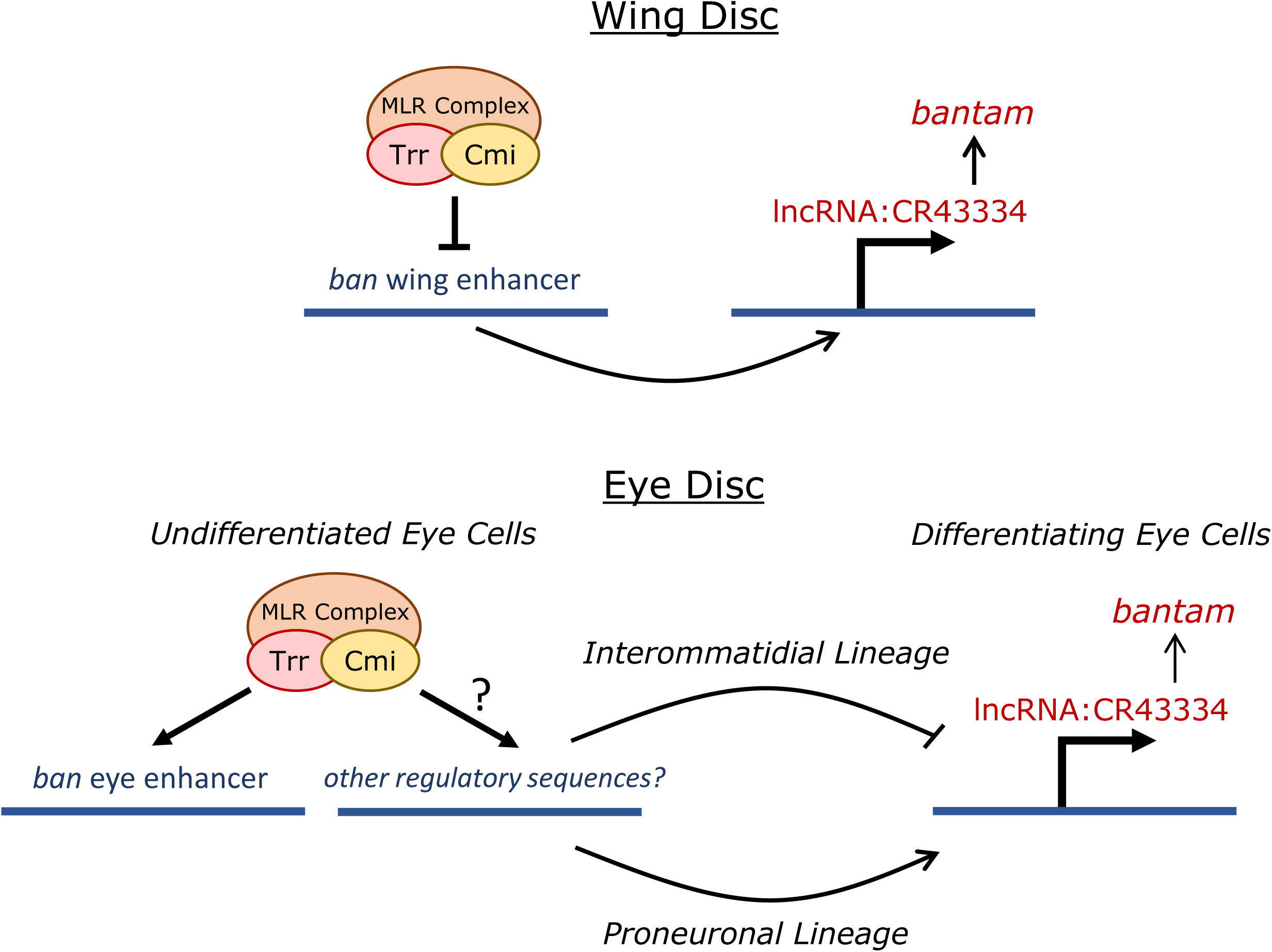
*Model of MLR functions in* bantam *enhancer control.* The Drosophila MLR COMPASS-like complex may either have positive or negative regulatory activity on the miRNA *bantam* depending on the context of cell fate. Tissue-specific *bantam* enhancers active in undifferentiated wing or eye tissue are bound and regulated by MLR. In the wing disc, MLR represses the activity of the *bantam* wing enhancer and downregulates *bantam* miRNA. In the eye disc, MLR modulates the expression pattern of the *bantam* eye enhancer in undifferentiated cells. After passage of the morphogenetic furrow and cell fate choice between proneuronal or interommatidial lineages, the previous regulatory activity of MLR leads to increased *bantam* expression in the proneuronal cells and decreased expression in the interommatidial.

We found that the MLR complex localizes to tissue-specific *bantam* enhancers during organ development, regulating the activity of those regions and the expression levels of the *bantam* miRNA. This regulatory activity is consequential for the development of these organs. We previously showed that the complex was vital for proper Dpp signaling during wing vein formation (Chauhan et al., 2013). In this report, we show that the complex is necessary for attenuating *bantam* expression in the wing via repression of a *cis*-acting distal *bantam* wing enhancer. *bantam* acts as a negative regulator of Dpp signaling through translation inhibition of Mad, a downstream effector of Dpp (Kane et al., 2018); thus, the vein patterning defects resulting from altered Dpp signaling upon reduction of MLR complex activity may be exacerbated by elevated *bantam* levels. These results reveal that proper expression control of *bantam* by the MLR complex is necessary for correct tissue patterning during wing development, likely through regulation of Dpp/Tgf-β signaling.

The RNAi knockdown of *Cmi* and *trr* in the developing eye tissue produces a complex phenotype, including reduction of eye tissue and pattern disruptions in the remaining ommatidia. This likely is the result of multiple altered transcriptional targets at various stages of compound eye development. The rough and shrunken phenotype is reminiscent of eye malformations caused by increased cell death or altered developmental signaling (Brennecke et al., 2003; Chao et al., 2004; Nolo et al., 2006), and *bantam* is recognized as both an inhibitor of apoptosis as well as a feedback regulator of multiple signaling pathways (Brennecke et al., 2003; Kane et al., 2018; Oh and Irvine, 2011; Wu et al., 2017; Zhang et al., 2013). While *bantam* overexpression exclusively in the differentiating ommatidia suppresses apoptosis and results in eye tissue overgrowth (Thompson and Cohen, 2006), we demonstrate here that overexpression of the miRNA throughout the eye pouch induces caspase activation in undifferentiated cells and phenocopies the rough and shrunken eyes resulting from reduced functions of the MLR complex. Massive cell death during development can result in reduced organ size, yet we demonstrate not only that the adult phenotype is not caused by increased apoptosis during development, but that caspase inhibition enhances the severity of the phenotype, potentially through suppression of non-apoptotic developmental caspase function (Kuranaga and Miura, 2007; Lamkanfi et al., 2007; Nakajima and Kuranaga, 2017). The fact that undifferentiated cell loss due to apoptosis does not affect the adult organ can be explained by compensatory proliferation of surviving neighbor cells to replace those lost or other caspase-related growth signals (Fan and Bergmann, 2008; James and Bryant, 1981; Shinoda et al., 2019; Song et al., 1997). These data not only further elucidate *bantam*’s roles in eye development, but also favor the hypothesis that alteration of developmental signaling rather than growth regulation underlies the adult eye defects we observe.

The importance of the MLR complex in contributing to either positive or negative regulation of a single transcriptional target depending on cell fate context is a novel observation with critical developmental consequence. This regulatory decision is likely not due to activity inherent to the complex itself, but rather influenced by the transcription factors that recruit the complex to specific developmental enhancers. Transcriptional control of *bantam* is orchestrated via complex signaling factors that regulate bantam enhancers (Peng et al., 2009; Slattery et al., 2013). For example, while Hippo effector Yki binds to and activates both the *bantam* eye and wing enhancers, binding partner Hth is necessary for regulating the eye enhancer and controlling *bantam* expression in the eye while separate partner Sd is necessary for regulating *bantam* expression in the wing through the wing enhancer (Peng et al., 2009; Slattery et al., 2013; Zhang et al., 2008). Multiple regulatory inputs are necessary for guiding proper spatiotemporal expression of the *bantam* miRNA in imaginal tissue, and our data suggests that the MLR complex plays a critical role in translating these inputs into regulatory decisions.

Through depletion of MLR complex activity in different regions of the eye imaginal disc, we determined that the complex is required in undifferentiated eye cells for proper regulation of *bantam* expression that occurs after furrow progression and the onset of differentiation into ommatidia, developmental stages separated by at least two rounds of mitosis (Baker, 2001; Cagan, 2009). Our eye disc studies correlate with *in vitro* data that the Kmt2D MLR complex is dispensable for maintaining gene transcription in murine ESCs, but is required for the transcriptional reprogramming that occurs during differentiation and dedifferentiation (Wang et al., 2016). Thus, our data not only verifies this role of MLR complexes through an *in vivo* developmental model but further demonstrates that this necessary preparatory activity may occur cell generations prior to the reprogramming event. We did not observe widespread changes to *bantam* levels in undifferentiated tissue upon *Cmi/trr* knockdown in larval eye discs, suggesting that MLR complex activity is not required for maintaining widespread *bantam* expression after the initial activation event and establishment of enhancer functions that likely occurred during embryogenesis (Brennecke et al., 2003; Rickels et al., 2017; Rickels and Shilatifard, 2018; Zraly et al., 2020). Thus, the MLR complex appears to be dispensable for the maintenance of previously active enhancers that drive widespread activity during development, but essential for the *de novo* programming of tissue-specific enhancers that occurs at critical developmental transitions, perhaps similar to the enhancer priming functions of Kmt2D in murine ESCs (Wang et al., 2016).

We found that the MLR complex has an unanticipated role in cell survival separate from *bantam* regulation. Knockdown of *Cmi* or *trr* in undifferentiated cells of the eye disc results in cell-autonomous elevation of executioner caspase (Dcp-1) signaling and positive TUNEL staining, indicative of caspase cascade and apoptotic cell death. Previous investigations in the Drosophila eye disc as well as human germinal B cells have reported that deletion of MLR complex methyltransferases both induces apoptosis and provides a proliferative advantage in undifferentiated cells, possibly through inhibition of differentiation (Kanda et al., 2013; Ortega-Molina et al., 2015). These results and ours suggest that MLR complexes play critical roles in multipotent cell maintenance and preparation for differentiation during cell fate transitions. While the apoptotic phenotype caused by *Cmi/trr* knockdown in the larval eye disc is not directly caused by dysregulated *bantam* expression, its sensitivity to *bantam* levels suggests the likely involvement of signaling pathway(s) active in that area and regulated by both the MLR complex and *bantam*. While many developmental pathways are necessary for proper eye development, the Notch signaling pathway is activated at the dorsal-ventral midline to promote proliferation and survival (Chao et al., 2004), and Notch signaling is dependent on both the MLR complex and bantam regulation (Becam et al., 2011; Dhar et al., 2018; Giaimo et al., 2017; Oswald et al., 2016; Wu et al., 2017).

Clearer comprehension of MLR complexes’ roles in enhancer regulation during normal development paves the way to better determine the mechanisms of disease associated with loss of complex function. The ability to regulate a single transcriptional target in opposite directions depending on tissue type and cell fate choice is critical for normal development. The miRNA *bantam* has previously been identified as tumorigenic in *Drosophila* cancer models (Sander and Herranz, 2019). Potential human counterparts of *bantam* have been identified, including *mir-450b* that contains close sequence similarity (Ibanez-Ventoso et al., 2008) and has been described as suppressing cancer cell proliferation and inducing protective differentiation (Sun et al., 2014; Zhao et al., 2014). The human *mir-130a* can act as either oncogene or tumor suppressor, impacts drug resistance (Zhang et al., 2017), and is functionally orthologous to *bantam* in its feedback regulation of Hippo pathway signaling (Shen et al., 2015). The discovery of the importance of miRNAs in both normal development and cancer together with emerging evidence for the fine-tuning of miRNA expression though MLR-dependent enhancer regulation, provides a strong rationale for the expansion of efforts to define the role of epigenetic gene control in the context of cell fate decisions.

## Acknowledgements

We thank Manuel O. Diaz and Richard Schultz for helpful discussions and comments on the manuscript, Jordan Beach and the SSOM Department of Cell and Molecular Physiology Specialized Imaging Resource Center for use of the Zeiss LSM 880 Airyscan microscope, and Michael Sertwetnyk for assistance with Drosophila genetic analyses. Fly stocks were obtained from the Bloomington Drosophila Stock Center (NIH P40OD018537) and from Stephen Cohen and Richard Mann. SEM analyses were performed with the help of Joseph Schluep and Jacob Ciszek at the Loyola University Chicago SEM facility, supported by a grant from the National Science Foundation-Major Research Instrumentation (MRI) Program [1726994]. The research was supported by grants from the National Science Foundation [MCB-1413331 and MCB-1716431 to A.K.D.].

## References

Andersen, E.C., Horvitz, H.R., 2007. Two C. elegans histone methyltransferases repress lin-3 EGF transcription to inhibit vulval development. Development 134, 2991–2999.

Ang, S.Y., Uebersohn, A., Spencer, C.I., Huang, Y., Lee, J.E., Ge, K., Bruneau, B.G., 2016. KMT2D regulates specific programs in heart development via histone H3 lysine 4 di-methylation. Development 143, 810–821.

Attisano, L., Wrana, J.L., 2013. Signal integration in TGF-beta, WNT, and Hippo pathways. F1000Prime Rep 5, 17.

Baker, N.E., 2001. Cell proliferation, survival, and death in the Drosophila eye. Seminars in Cell & Developmental Biology 12, 499–507.

Becam, I., Rafel, N., Hong, X., Cohen, S.M., Milan, M., 2011. Notch-mediated repression of bantam miRNA contributes to boundary formation in the Drosophila wing. Development 138, 3781–3789.

Boulan, L., Martin, D., Milan, M., 2013. bantam miRNA promotes systemic growth by connecting insulin signaling and ecdysone production. Curr Biol 23, 473–478.

Brachmann, C.B., Cagan, R.L., 2003. Patterning the fly eye: the role of apoptosis. Trends Genet 19, 91–96.

Brand, A.H., Manoukian, A.S., Perrimon, N., 1994. Ectopic expression in Drosophila. Methods Cell Biol 44, 635–654.

Brennecke, J., Hipfner, D.R., Stark, A., Russell, R.B., Cohen, S.M., 2003. bantam encodes a developmentally regulated microRNA that controls cell proliferation and regulates the proapoptotic gene hid in Drosophila. Cell 113, 25–36.

Cagan, R., 2009. Principles of Drosophila eye differentiation. Curr Top Dev Biol 89, 115–135.

Chao, J.L., Tsai, Y.C., Chiu, S.J., Sun, Y.H., 2004. Localized Notch signal acts through eyg and upd to promote global growth in Drosophila eye. Development 131, 3839–3847.

Chauhan, C., Zraly, C.B., Dingwall, A.K., 2013. The Drosophila COMPASS-like Cmi-Trr coactivator complex regulates dpp/BMP signaling in pattern formation. Developmental Biology 380, 185–198.

Chauhan, C., Zraly, C.B., Parilla, M., Diaz, M.O., Dingwall, A.K., 2012. Histone recognition and nuclear receptor co-activator functions of Drosophila Cara Mitad, a homolog of the N-terminal portion of mammalian MLL2 and MLL3. Development 139, 1997–2008.

Dhar, S.S., Zhao, D., Lin, T., Gu, B., Pal, K., Wu, S.J., Alam, H., Lv, J., Yun, K., Gopalakrishnan, V., Flores, E.R., Northcott, P.A., Rajaram, V., Li, W., Shilatifard, A., Sillitoe, R.V., Chen, K., Lee, M.G., 2018. MLL4 Is Required to Maintain Broad H3K4me3 Peaks and Super-Enhancers at Tumor Suppressor Genes. Mol Cell 70, 825–841 e826.

Doumpas, N., Ruiz-Romero, M., Blanco, E., Edgar, B., Corominas, M., Teleman, A.A., 2013. Brk regulates wing disc growth in part via repression of Myc expression. EMBO Rep 14, 261–268.

Fagan, R.J., Dingwall, A.K., 2019. COMPASS Ascending: Emerging clues regarding the roles of MLL3/KMT2C and MLL2/KMT2D proteins in cancer. Cancer Lett 458, 56–65.

Fan, Y., Bergmann, A., 2008. Distinct mechanisms of apoptosis-induced compensatory proliferation in proliferating and differentiating tissues in the Drosophila eye. Dev Cell 14, 399–410.

Ford, D.J., Dingwall, A.K., 2015. The cancer COMPASS: navigating the functions of MLL complexes in cancer. Cancer Genet 208, 178–191.

Gerlach, S.U., Sander, M., Song, S., Herranz, H., 2019. The miRNA bantam regulates growth and tumorigenesis by repressing the cell cycle regulator tribbles. Life Sci Alliance 2.

Giaimo, B.D., Oswald, F., Borggrefe, T., 2017. Dynamic chromatin regulation at Notch target genes. Transcription 8, 61–66.

Grether, M.E., Abrams, J.M., Agapite, J., White, K., Steller, H., 1995. The head involution defective gene of Drosophila melanogaster functions in programmed cell death. Genes Dev 9, 1694–1708.

Herranz, H., Hong, X., Cohen, S.M., 2012. Mutual repression by bantam miRNA and Capicua links the EGFR/MAPK and Hippo pathways in growth control. Curr Biol 22, 651–657.

Herranz, H., Hong, X., Perez, L., Ferreira, A., Olivieri, D., Cohen, S.M., Milan, M., 2010. The miRNA machinery targets Mei-P26 and regulates Myc protein levels in the Drosophila wing. EMBO J 29, 1688–1698.

Herz, H.M., Mohan, M., Garruss, A.S., Liang, K., Takahashi, Y.H., Mickey, K., Voets, O., Verrijzer, C.P., Shilatifard, A., 2012. Enhancer-associated H3K4 monomethylation by Trithorax-related, the Drosophila homolog of mammalian Mll3/Mll4. Genes Dev 26, 2604–2620.

Hipfner, D.R., Weigmann, K., Cohen, S.M., 2002. The bantam gene regulates Drosophila growth. Genetics 161, 1527–1537.

Hu, D., Garruss, A.S., Gao, X., Morgan, M.A., Cook, M., Smith, E.R., Shilatifard, A., 2013. The Mll2 branch of the COMPASS family regulates bivalent promoters in mouse embryonic stem cells. Nat Struct Mol Biol 20, 1093–1097.

Ibanez-Ventoso, C., Vora, M., Driscoll, M., 2008. Sequence relationships among C. elegans, D. melanogaster and human microRNAs highlight the extensive conservation of microRNAs in biology. PLoS One 3, e2818.

Issaeva, I., Zonis, Y., Rozovskaia, T., Orlovsky, K., Croce, C.M., Nakamura, T., Mazo, A., Eisenbach, L., Canaani, E., 2007. Knockdown of ALR (MLL2) reveals ALR target genes and leads to alterations in cell adhesion and growth. Mol Cell Biol 27, 1889–1903.

James, A.A., Bryant, P.J., 1981. A quantitative study of cell death and mitotic inhibition in gamma-irradiated imaginal wing discs of Drosophila melanogaster. Radiat Res 87, 552–564.

Jordan-Alvarez, S., Santana, E., Casas-Tinto, S., Acebes, A., Ferrus, A., 2017. The equilibrium between antagonistic signaling pathways determines the number of synapses in Drosophila. PLoS One 12, e0184238.

Kanda, H., Nguyen, A., Chen, L., Okano, H., Hariharan, I.K., 2013. The Drosophila ortholog of MLL3 and MLL4, trithorax related, functions as a negative regulator of tissue growth. Mol Cell Biol 33, 1702–1710.

Kane, N.S., Vora, M., Padgett, R.W., Li, Y., 2018. bantam microRNA is a negative regulator of the Drosophila decapentaplegic pathway. Fly (Austin) 12, 105–117.

Kleefstra, T., Kramer, J.M., Neveling, K., Willemsen, M.H., Koemans, T.S., Vissers, L.E., Wissink-Lindhout, W., Fenckova, M., van den Akker, W.M., Kasri, N.N., Nillesen, W.M., Prescott, T., Clark, R.D., Devriendt, K., van Reeuwijk, J., de Brouwer, A.P., Gilissen, C., Zhou, H., Brunner, H.G., Veltman, J.A., Schenck, A., van Bokhoven, H., 2012. Disruption of an EHMT1-associated chromatin-modification module causes intellectual disability. American journal of human genetics 91, 73–82.

Kuranaga, E., Miura, M., 2007. Nonapoptotic functions of caspases: caspases as regulatory molecules for immunity and cell-fate determination. Trends Cell Biol 17, 135–144.

Lai, B., Lee, J.E., Jang, Y., Wang, L., Peng, W., Ge, K., 2017. MLL3/MLL4 are required for CBP/p300 binding on enhancers and super-enhancer formation in brown adipogenesis. Nucleic Acids Res 45, 6388–6403.

Lamkanfi, M., Festjens, N., Declercq, W., Vanden Berghe, T., Vandenabeele, P., 2007. Caspases in cell survival, proliferation and differentiation. Cell Death Differ 14, 44–55.

Lee, J.E., Wang, C., Xu, S., Cho, Y.W., Wang, L., Feng, X., Baldridge, A., Sartorelli, V., Zhuang, L., Peng, W., Ge, K., 2013. H3K4 mono- and di-methyltransferase MLL4 is required for enhancer activation during cell differentiation. Elife 2, e01503.

Levine, M., 2010. Transcriptional enhancers in animal development and evolution. Curr Biol 20, R754–763.

Li, Y., Padgett, R.W., 2012. bantam is required for optic lobe development and glial cell proliferation. PLoS One 7, e32910.

Ma, X., Wang, H., Ji, J., Xu, W., Sun, Y., Li, W., Zhang, X., Chen, J., Xue, L., 2017. Hippo signaling promotes JNK-dependent cell migration. Proc Natl Acad Sci U S A 114, 1934–1939.

Martin, F.A., Perez-Garijo, A., Moreno, E., Morata, G., 2004. The brinker gradient controls wing growth in Drosophila. Development 131, 4921–4930.

Nakajima, Y.I., Kuranaga, E., 2017. Caspase-dependent non-apoptotic processes in development. Cell Death Differ 24, 1422–1430.

Nan, Z., Yang, W., Lyu, J., Wang, F., Deng, Q., Xi, Y., Yang, X., Ge, W., 2019. Drosophila Hcf regulates the Hippo signaling pathway via association with the histone H3K4 methyltransferase Trr. Biochem J 476, 759–768.

Ng, S.B., Bigham, A.W., Buckingham, K.J., Hannibal, M.C., McMillin, M.J., Gildersleeve, H.I., Beck, A.E., Tabor, H.K., Cooper, G.M., Mefford, H.C., Lee, C., Turner, E.H., Smith, J.D., Rieder, M.J., Yoshiura, K., Matsumoto, N., Ohta, T., Niikawa, N., Nickerson, D.A., Bamshad, M.J., Shendure, J., 2010. Exome sequencing identifies MLL2 mutations as a cause of Kabuki syndrome. Nat Genet 42, 790–793.

Nolo, R., Morrison, C.M., Tao, C., Zhang, X., Halder, G., 2006. The bantam microRNA is a target of the hippo tumor-suppressor pathway. Current biology : CB 16, 1895–1904.

Oh, H., Irvine, K.D., 2011. Cooperative regulation of growth by Yorkie and Mad through bantam. Developmental cell 20, 109–122.

Oh, H., Slattery, M., Ma, L., Crofts, A., White, K.P., Mann, R.S., Irvine, K.D., 2013. Genome-wide association of Yorkie with chromatin and chromatin-remodeling complexes. Cell Rep 3, 309–318.

Oh, H., Slattery, M., Ma, L., White, K.P., Mann, R.S., Irvine, K.D., 2014. Yorkie promotes transcription by recruiting a histone methyltransferase complex. Cell Rep 8, 449–459.

Ortega-Molina, A., Boss, I.W., Canela, A., Pan, H., Jiang, Y., Zhao, C., Jiang, M., Hu, D., Agirre, X., Niesvizky, I., Lee, J.E., Chen, H.T., Ennishi, D., Scott, D.W., Mottok, A., Hother, C., Liu, S., Cao, X.J., Tam, W., Shaknovich, R., Garcia, B.A., Gascoyne, R.D., Ge, K., Shilatifard, A., Elemento, O., Nussenzweig, A., Melnick, A.M., Wendel, H.G., 2015. The histone lysine methyltransferase KMT2D sustains a gene expression program that represses B cell lymphoma development. Nat Med 21, 1199–1208.

Oswald, F., Rodriguez, P., Giaimo, B.D., Antonello, Z.A., Mira, L., Mittler, G., Thiel, V.N., Collins, K.J., Tabaja, N., Cizelsky, W., Rothe, M., Kuhl, S.J., Kuhl, M., Ferrante, F., Hein, K., Kovall, R.A., Dominguez, M., Borggrefe, T., 2016. A phospho-dependent mechanism involving NCoR and KMT2D controls a permissive chromatin state at Notch target genes. Nucleic Acids Res.

Peng, H.W., Slattery, M., Mann, R.S., 2009. Transcription factor choice in the Hippo signaling pathway: homothorax and yorkie regulation of the microRNA bantam in the progenitor domain of the Drosophila eye imaginal disc. Genes Dev 23, 2307–2319.

Qing, Y., Yin, F., Wang, W., Zheng, Y., Guo, P., Schozer, F., Deng, H., Pan, D., 2014. The Hippo effector Yorkie activates transcription by interacting with a histone methyltransferase complex through Ncoa6. Elife 3.

Qu, Z., Bendena, W.G., Nong, W., Siggens, K.W., Noriega, F.G., Kai, Z.P., Zang, Y.Y., Koon, A.C., Chan, H.Y.E., Chan, T.F., Chu, K.H., Lam, H.M., Akam, M., Tobe, S.S., Lam Hui, J.H., 2017. MicroRNAs regulate the sesquiterpenoid hormonal pathway in Drosophila and other arthropods. Proc Biol Sci 284.

Rickels, R., Herz, H.M., Sze, C.C., Cao, K., Morgan, M.A., Collings, C.K., Gause, M., Takahashi, Y.H., Wang, L., Rendleman, E.J., Marshall, S.A., Krueger, A., Bartom, E.T., Piunti, A., Smith, E.R., Abshiru, N.A., Kelleher, N.L., Dorsett, D., Shilatifard, A., 2017. Histone H3K4 monomethylation catalyzed by Trr and mammalian COMPASS-like proteins at enhancers is dispensable for development and viability. Nat Genet 49, 1647–1653.

Rickels, R., Shilatifard, A., 2018. Enhancer Logic and Mechanics in Development and Disease. Trends Cell Biol 28, 608–630.

Sander, M., Herranz, H., 2019. MicroRNAs in Drosophila Cancer Models. Adv Exp Med Biol 1167, 157–173.

Schindelin, J., Arganda-Carreras, I., Frise, E., Kaynig, V., Longair, M., Pietzsch, T., Preibisch, S., Rueden, C., Saalfeld, S., Schmid, B., Tinevez, J.Y., White, D.J., Hartenstein, V., Eliceiri, K., Tomancak, P., Cardona, A., 2012. Fiji: an open-source platform for biological-image analysis. Nat Methods 9, 676–682.

Sedkov, Y., Benes, J.J., Berger, J.R., Riker, K.M., Tillib, S., Jones, R.S., Mazo, A., 1999. Molecular genetic analysis of the Drosophila trithorax-related gene which encodes a novel SET domain protein. Mech Dev 82, 171–179.

Sedkov, Y., Cho, E., Petruk, S., Cherbas, L., Smith, S.T., Jones, R.S., Cherbas, P., Canaani, E., Jaynes, J.B., Mazo, A., 2003. Methylation at lysine 4 of histone H3 in ecdysone-dependent development of Drosophila. Nature 426, 78–83.

Shen, S., Guo, X., Yan, H., Lu, Y., Ji, X., Li, L., Liang, T., Zhou, D., Feng, X.H., Zhao, J.C., Yu, J., Gong, X.G., Zhang, L., Zhao, B., 2015. A miR-130a-YAP positive feedback loop promotes organ size and tumorigenesis. Cell Res 25, 997–1012.

Shilatifard, A., 2012. The COMPASS family of histone H3K4 methylases: mechanisms of regulation in development and disease pathogenesis. Annu Rev Biochem 81, 65–95.

Shinoda, N., Hanawa, N., Chihara, T., Koto, A., Miura, M., 2019. Dronc-independent basal executioner caspase activity sustains Drosophila imaginal tissue growth. Proc Natl Acad Sci U S A 116, 20539–20544.

Slattery, M., Voutev, R., Ma, L., Negre, N., White, K.P., Mann, R.S., 2013. Divergent transcriptional regulatory logic at the intersection of tissue growth and developmental patterning. PLoS Genet 9, e1003753.

Song, Z., Guan, B., Bergman, A., Nicholson, D.W., Thornberry, N.A., Peterson, E.P., Steller, H., 2000. Biochemical and genetic interactions between Drosophila caspases and the proapoptotic genes rpr, hid, and grim. Mol Cell Biol 20, 2907–2914.

Song, Z., McCall, K., Steller, H., 1997. DCP-1, a Drosophila cell death protease essential for development. Science 275, 536–540.

Sun, M.M., Li, J.F., Guo, L.L., Xiao, H.T., Dong, L., Wang, F., Huang, F.B., Cao, D., Qin, T., Yin, X.H., Li, J.M., Wang, S.L., 2014. TGF-beta1 suppression of microRNA-450b-5p expression: a novel mechanism for blocking myogenic differentiation of rhabdomyosarcoma. Oncogene 33, 2075–2086.

Tanaka-Matakatsu, M., Xu, J., Cheng, L., Du, W., 2009. Regulation of apoptosis of rbf mutant cells during Drosophila development. Dev Biol 326, 347–356.

Thompson, B.J., Cohen, S.M., 2006. The Hippo pathway regulates the bantam microRNA to control cell proliferation and apoptosis in Drosophila. Cell 126, 767–774.

Van Laarhoven, P.M., Neitzel, L.R., Quintana, A.M., Geiger, E.A., Zackai, E.H., Clouthier, D.E., Artinger, K.B., Ming, J.E., Shaikh, T.H., 2015. Kabuki syndrome genes KMT2D and KDM6A: functional analyses demonstrate critical roles in craniofacial, heart and brain development. Hum Mol Genet 24, 4443–4453.

Wang, C., Lee, J.E., Lai, B., Macfarlan, T.S., Xu, S., Zhuang, L., Liu, C., Peng, W., Ge, K., 2016. Enhancer priming by H3K4 methyltransferase MLL4 controls cell fate transition. Proc Natl Acad Sci U S A 113, 11871–11876.

Wang, L., Zhao, Z., Ozark, P.A., Fantini, D., Marshall, S.A., Rendleman, E.J., Cozzolino, K.A., Louis, N., He, X., Morgan, M.A., Takahashi, Y.H., Collings, C.K., Smith, E.R., Ntziachristos, P., Savas, J.N., Zou, L., Hashizume, R., Meeks, J.J., Shilatifard, A., 2018. Resetting the epigenetic balance of Polycomb and COMPASS function at enhancers for cancer therapy. Nat Med 24, 758–769.

Weng, R., Cohen, S.M., 2015. Control of Drosophila Type I and Type II central brain neuroblast proliferation by bantam microRNA. Development 142, 3713–3720.

Wolff, T., 2011. Preparation of Drosophila eye specimens for scanning electron microscopy. Cold Spring Harb Protoc 2011, 1383–1385.

Wu, Y.C., Lee, K.S., Song, Y., Gehrke, S., Lu, B., 2017. The bantam microRNA acts through Numb to exert cell growth control and feedback regulation of Notch in tumor-forming stem cells in the Drosophila brain. PLoS Genet 13, e1006785.

Zhang, H.D., Jiang, L.H., Sun, D.W., Li, J., Ji, Z.L., 2017. The role of miR-130a in cancer. Breast Cancer 24, 521–527.

Zhang, L., Ren, F., Zhang, Q., Chen, Y., Wang, B., Jiang, J., 2008. The TEAD/TEF family of transcription factor Scalloped mediates Hippo signaling in organ size control. Dev Cell 14, 377–387.

Zhang, X., Luo, D., Pflugfelder, G.O., Shen, J., 2013. Dpp signaling inhibits proliferation in the Drosophila wing by Omb-dependent regional control of bantam. Development 140, 2917–2922.

Zhao, Z., Li, R., Sha, S., Wang, Q., Mao, W., Liu, T., 2014. Targeting HER3 with miR-450b-3p suppresses breast cancer cells proliferation. Cancer Biol Ther 15, 1404–1412.

Zoog, S.J., Bertin, J., Friesen, P.D., 1999. Caspase inhibition by baculovirus P35 requires interaction between the reactive site loop and the beta-sheet core. J Biol Chem 274, 25995–26002.

Zraly, C.B., Zakkar, A., Perez, J.H., Ng, J., White, K.P., Slattery, M., Dingwall, A.K., 2020. The Drosophila MLR COMPASS complex is essential for programming cis-regulatory information and maintaining epigenetic memory during development. Nucleic Acids Res, 1–20.

